# TEclass2: Classification of transposable elements using Transformers

**DOI:** 10.1101/2023.10.13.562246

**Authors:** Lucas Bickmann, Matias Rodriguez, Xiaoyi Jiang, Wojciech Makalowski

## Abstract

**Motivation:** Transposable elements (TEs) are interspersed repetitive sequences that are major constituents of most eukaryotic genomes and are crucial for genome evolution. Despite the existence of multiple tools for their classification and annotation, none of them can achieve completely reliable results making it a challenge for genomic studies. In this work, we introduce TEclass2, a new software that uses a deep learning approach based upon a linear Transformer architecture with a k-mer tokenizer and further adaptations to handle DNA sequences. This software has an easy configuration that allows training models on new datasets and the classification of TE models providing multiple metrics for a reliable evaluation of the results.

**Results:** This work shows a successful adaptation of deep learning with Transformers for the classification of TE models from consensus sequences, and these results lay a foundation for novel methodologies in bioinformatics. We provide a tool for the training of models and the classification of consensus sequences from TE models on custom data and a web page interface with a pre-trained dataset based on curated and non-curated TE libraries allowing a fast and simple classification of TEs.

**Availability:** https://bioinformatics.uni-muenster.de/tools/teclass2/index.pl

**Supplementary information:** Supplementary data are available at *Bioinformatics* online.

## 1 Introduction

Transposable elements (TEs) are mobile genetic elements of possible viral origin which are present in great numbers in the majority of eukaryotic genomes. TEs constitute up to 46% of the human genome (Hoyt *et al*., 2022) and a bulk of many plant genomes such as wheat and maize where TEs comprise 85% of the genome (Wicker *et al*., 2018) (Schnable *et al*., 2009). TEs are major drivers of genome evolution as they can facilitate chromosome rearrangements by acting as recombination hotspots, they can also provide mechanisms for genomic shuffling (Moran *et al*., 1999), and contribute in genome expansion (Piegu *et al*., 2006) (Ungerer *et al*., 2006), completely altering the whole architecture of a genome. TEs can also modify gene expression by disrupting regulatory sequences, impairing genes, and promoting the emergence of new sequences (Makałowski, 2000).

Despite the abundance of TEs in most genomes, the identification of TEs is still challenging and time-consuming, as TEs can be extremely diverse in their DNA sequence due to varying motifs, lengths and structures. It is also common to find very old TE families where the majority of TE sequences are inactive due to an accumulation of mutations and fragmentation. This means that different copies of the same TE can be very different from each other, making the identification of decayed copies a very difficult task (Rodriguez and Makałowski, 2022).

For the identification of TEs there are two main strategies, one is to perform a homology search on the genome using TE-sequences’ libraries. Another way of finding TEs, which is particularly useful in non-model organisms or when the aim is to study or discover new TEs is to use a ’de novo’ approach, which initially does not rely in previous information. In a ’de novo’ approach interspersed repetitive sequences from the genome are identified and used to build consensus models of candidate TEs, which are compared to previously recorded ones from TE libraries and then are classified and annotated.

Multiple approaches can be used for the classification of TEs depending on structural features, transposition mechanisms, enzymatic machinery or the evolutionary context (Piégu *et al*., 2015). The first proposal for classifying TEs (Finnegan, 1989) divided them into two classes based on their mechanism of transposition. The TEs that use the reverse transcription of an RNA intermediate are called class I elements or retrotransposons, while the TEs that transpose using a DNA intermediate are called class II elements or DNA transposons. Class I TEs were further divided into two categories, LTR-retrotransposons and non-LTR retrotransposons. LTR-retrotransposons are similar to retroviruses, with open reading frames flanked by long terminal repeats (LTR), whereas non-LTR retrotransposons have no LTRs or very short terminal repeats and a poly-A tail.

This initial classification did not consider non autonomous elements, such as SINEs, and has been revisited and updated multiple times. The Repbase (Jurka *et al*., 2005) (Kapitonov and Jurka, 2008) and Wicker (Wicker *et al*., 2007) classifications are probably the best well-known proposals. Both classifications added a number of different TE subdivisions which further classify the original two TE classes taking in consideration sequence similarities, sequence structure, and the number of ORFs and enzymes coded. The Repbase classification divides TEs in three main classes: DNA transposons, LTR retrotransposons and non-LTR retrotransposons; each one of these classes has further subdivisions with the whole library having 65 superfamilies or clades (Bao *et al*., 2015). In the Wicker proposal for classifying TEs, the Class I elements are divided into five orders, namely LTR, DIRS, PLE, LINEs, and SINEs. Class II elements are divided in two subclasses: subclass 1 are TEs that use a cut and paste mechanism and consist of TIR and Crypton TEs, while subclass 2 are TEs with a copy and paste mechanism and consist of Helitron and Maverick TEs (Wicker *et al*., 2007).

The necessity to identify and classify these huge parts of the genome pushed for the development of pipelines that allow a semiautomated discovery and annotation of TEs. Most commonly used tools classify TEs based on sequence similarity by doing similarity comparisons of consensus sequences to databases of TEs. Among popular software packages, RepeatModeler uses the DFAM database for TE classification and REPET uses its own database RepetDB with a classification hierarchy similar to Repbase. There are alternative strategies to sequence homology comparisons, such as the software PASTEC (Hoede *et al*., 2014) which is part of the REPET pipeline and uses HMM profiles to identify TEs.

Another alternative strategy is to use machine learning, a field that has shown promising results in bioinformatics (Wu and Zhao, 2019). Its application has become more reliable and feasible in recent years as the amount of genomic data being produced has increased significantly and there have been multiple advances in deep learning algorithms which have improved the speed and accuracy of this approach (Lan *et al*., 2018) (Li *et al*., 2022). Machine learning techniques can automatically identify patterns and extract information from data to solve problems of classification of strings of text such as the DNA sequences.

The first software to use machine learning for classifying TEs was TEclass (Abrusán *et al*., 2009), which implemented support vector machines (SVM) to classify unknown TEs using the oligomer frequencies of tetramers and pentamers found in repeat elements. TEs are classified into their main categories as DNA transposons, LTRs and non-LTR repeats, and as LINEs or SINEs. These classifiers were built using TE sequences from Repbase and follow a binary classification in each step, considering initially if TEs are DNA transposons or Retrotransposons, in the next step LTRs versus non-LTRs for retroelements, and then LINEs versus SINEs for non-LTR elements (Abrusán *et al*., 2009).

In recent years, machine-learning methods that use deep learning strategies based on neural networks have enabled the development of new tools. For the classification of TEs into their respective orders or superfamilies, some software such as DeepTE (Yan *et al*., 2020) and TERL (da Cruz *et al*., 2021), are based on the machine learning architecture of convolutional neural networks (CNNs). Among the multiple tools for classifying TEs, some are specific to a certain order of TEs. For example, TIR-Learner (Su *et al*., 2019) is used to classify

TIR TEs using neural networks, k-nearest neighbor, random forest, and Adaboost. TE-Learner (Schietgat *et al*., 2018) classifies LTR TEs using a random forest approach, while Inpactor2 (Orozco-Arias *et al*., 2023) classifies LTR-retrotransposons from plant genomes using an architecture based on CNNs. In the TEclass2 presented here, we use the deep learning model Transformers (Vaswani *et al*., 2017) that was developed for natural language processing (NLP) and computer vision. It performs exceptionally well in these fields even replacing other models such as recurrent neural networks (RNNs) due to improved performance (Wolf *et al*., 2020). Transformers have the ability to process an entire input all at once by using an attention mechanism that provides a context for any position in a string, allowing parallelization and reducing the time used for training datasets (Vaswani *et al*., 2017).

Some other works also introduced the usage of Transformers into the field of bioinformatics, such as prediction of promoters, splice sites, and transcription factor binding sites by DNABERT (Ji *et al*., 2021) based in the Transformer language model used for NLP, called BERT (Devlin *et al*., 2019). Similar approaches can be used for bacterial promoter classification, viral genome identification, and mRNA degradation properties of COVID-19 vaccine candidates (He *et al*., 2021).

The proposed transformer-based models show state-of-the-art performance in their respective fields, are based upon the vanilla Transformer or BERT and their runtime and space-complexity is O(*n*^2^), performing usually up to 512 tokens. This limitation has been attempted to be tackled with linear transformer models, such as the Linformer (Wang *et al*., 2020) and Longformer (Beltagy *et al*., 2020). Depending on the available memory, they can process up to 4096 tokens or even longer, however, these architectures are not yet adapted to non-separated strings such as DNA sequences.

The work presented here introduces a new architecture based on the Longformer model (Beltagy *et al*., 2020) for the classification of selected TEs sequences, including various sequence specific augmentations, a k-mer specialized tokenizer, and implementing sliding window dilation. TEclass2 is an all-in-one classifier that can be used to rapidly predict TE orders and superfamilies using TE models built upon the novel Transformer architecture. The software is accessible through a web interface where it offers the possibility to classify sequences into sixteen superfamilies from the Wicker classification system (Wicker *et al*., 2007). Alternatively, users can download the source code to train and build their own classification models.

## 2 Methods

### 2.1 Transformer

A Transformer is a deep learning model that uses the mechanism of self-attention originally designed to handle sequential input, especially natural language, and is vastly improved in its performance when compared to its predecessors, the Recurrent Neural Networks (RNNs) (Rumelhart *et al*., 1986) and Long Short-Term Memory (LSTM) (Hochreiter and Schmidhuber, 1997). Instead of using the input as a sequence and processing each word in order as RNNs do, they process the whole sentence concurrently, allowing parallel computation and improving long-term dependencies. Originally transformers were designed using sequence transformation with a Seq2Seq model for performing tasks like translation and summarization of text. This model has an encoder that captures the input and passes it to a decoder that produces an output (Sutskever *et al*., 2014).

In TEclass2, for the classification of TE DNA sequences, we use only the encoder-block, followed by a classification head as in a linear layer (Figure 1). The transformer model is divided into several layers which have multiple attention heads (Luong *et al*., 2015) and each one that can learn independently relevant parts of the input sequence.

**Fig. 1.**
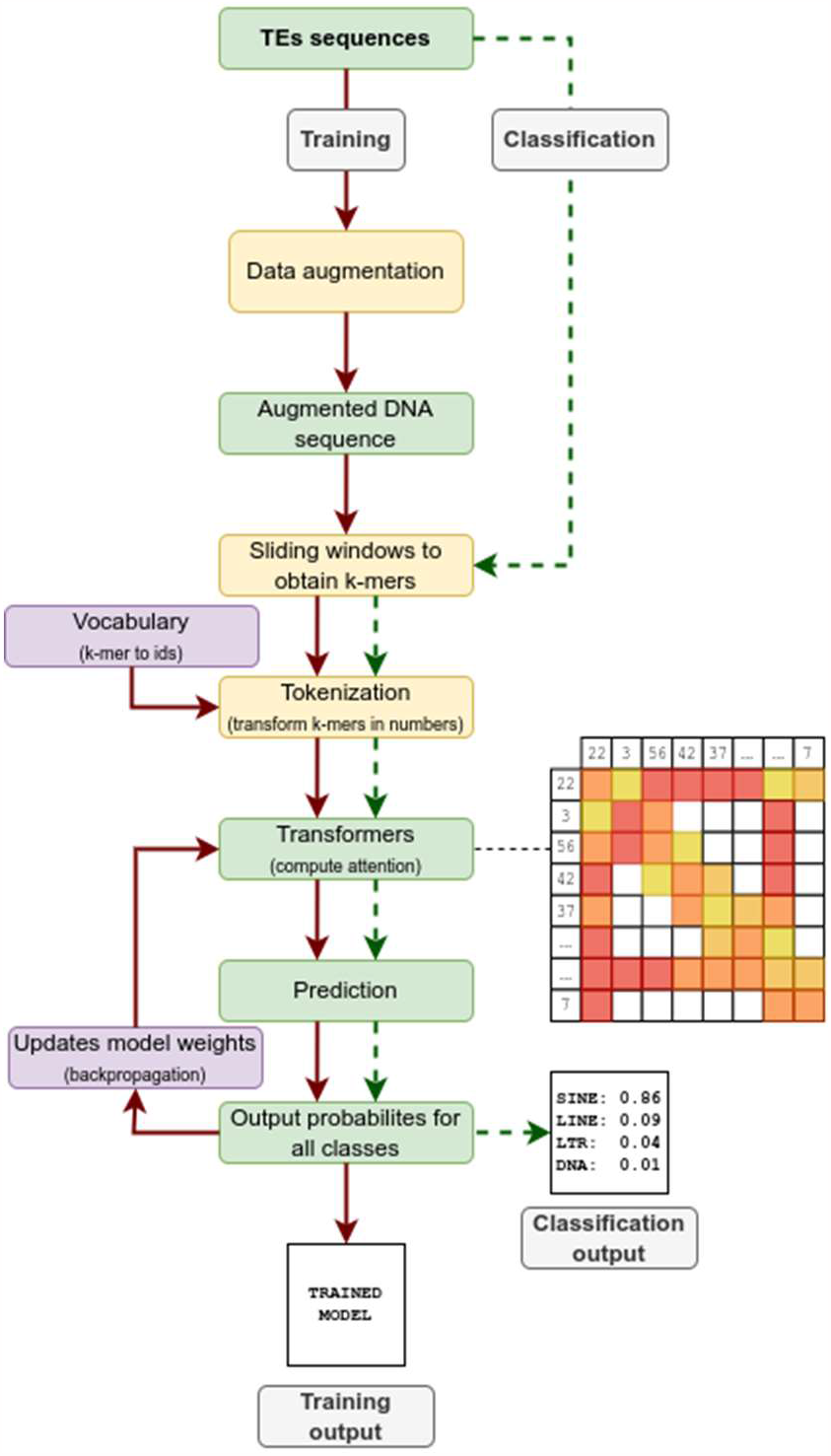
Flowchart succinctly describing how TEclass2 works both in the training of dataset to produce Transformer models (red arrows) and the use of these models to classify TE sequences (dotted green arrows)

The first layer’s query is the encoded query or tokens and the following layers are based upon previously computed attention matrices as queries. In order to take into consideration the order of the input sequence, we added positional encoding. Due to memory constraints transformers can process up to 512 tokens and larger inputs are either truncated or processed in chunks. For the latter the classification is then computed as the median of the output from each of the processed chunks. To solve this problem, different methods have been proposed such as the Longformer method which tackles this with reduced self-attention masks of local attentions and by adding some global-attentions (Beltagy *et al*., 2020).

A way to increase variants in the data during training is to us data augmentation and forcing the machine learning model to adapt to more generalized features (Shorten and Khoshgoftaar, 2019). To apply an on-the-fly sequence augmentation during the training step, we developed a variety of biologically motivated data augmentations (Table 1).

**Table 1.**
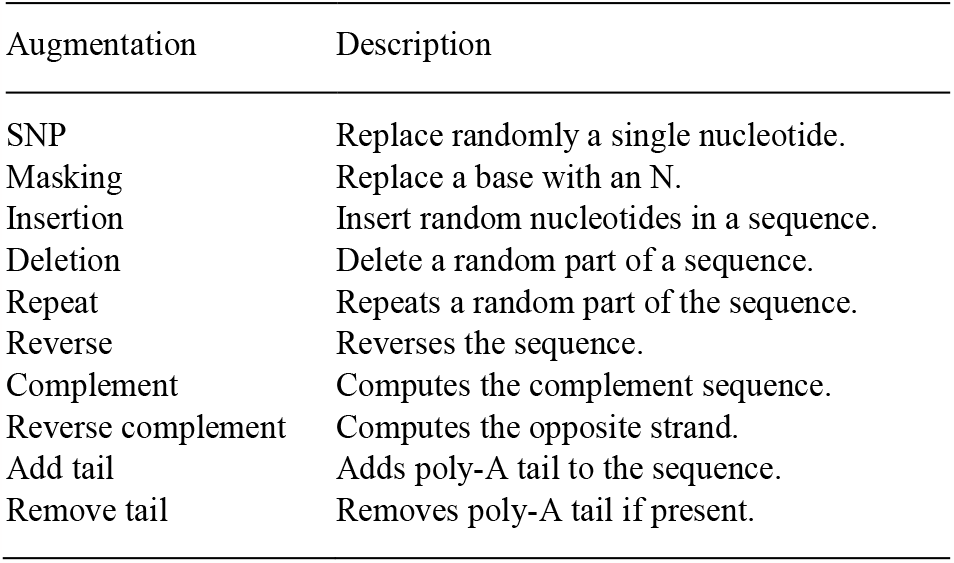
List of data augmentation used in DNA sequences for the data-training step in TEclass2.

Depending on the applied augmentations, the DNA sequence may also differ in length. The only transformation that introduces ambiguity is the introduction of Ns in the sequence and in this case the transformer should learn that these characters do not represent relevant information. The original transformer does not handle information about the position of tokens due to the transformers parallel processing of the data, therefore the relationship between tokens has to be provided, as these distances provide important spatial information. Sinusoidal positional encoding is used for this as it has the advantage of providing information that is independent from the length of the input and each value ranges between zero and one (Shaw *et al*., 2018).

In TEclass2 the DNA sequences used as an input undergo a process known as tokenization, in which the input strings are split into smaller parts called tokens which are assigned identifiers based on a created vocabulary. Tokenizers can also be embedded into a vector space and used as a form of compression and dictionary coding. We used k-mer tokenization with a sliding window approach dividing each DNA sequence into k-mers, whereas the general length of the embedded string does not decrease when using a different window size. A larger window size dilates the steps between the k-mers, which increases the coverage of the input sequence but reduces the overlap and dependencies between them, as they also have the advantage of generating a fixed number of words. In our software we used k-mers with a length of five with a sliding step of two nucleotides, the number of DNA 5-mers including unknown nucleotides as N gives a rather small vocabulary of 3125 tokens. With this software it is possible to use k-mers of different length but it’s important to consider that using the shorter ones impedes the encoding of different semantic representations, and using the larger ones increases the size of the vocabulary manifold degrading the performance.

In order to allow long DNA sequences we introduced a dynamically scaling sliding step of minimal two nucleotides to maximize the coverage of the input sequence, while trying to retain overlapping tokens. We compute it as:

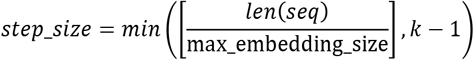

The attention values are traced back to each token and using linear interpolation these values are scaled back. This reduces cross dependencies between local tokens and increases long-range dependencies, which are useful to classify long TE sequences.

The models tested were trained with weighted cross entropy (WCE), as the datasets are unbalanced. The Softmax function (Goodfellow *et al*., 2016) is used to compute the probabilities for predicting the TE classes and to classify the performance of each trained model we used different metrics during evaluation and testing. This includes the train-and evaluation-loss, class-wise precision, recall, F1-score, and global accuracy on the test-dataset; also a macro average and weighted average were computed for precision, recall, and F1-score.

### 2.2 Workflow

The input data for training and building a model with TEclass2 is a FASTA file with the TE models, where the header of each sequences contains the name and the class of TE. For the training of a model, the workflow (Figure 1) starts by building a database of labeled TE sequences, which is divided into training, evaluation and testing datasets. The configuration of the database is read from the configuration file, where the expected family names and files paths must be specified.

After the database has been built, the training step, used to train the model, can begin utilizing parameters from the configuration file such as the number of epochs, the size of the layers for the neural network, size of the embeddings, window dilation, and many others. The training step uses the data augmentation properties specified in Table 1, which increase the variability of the input sequences, then, the TE sequences are scanned using a sliding window approach.

Next, each k-mer is tokenized and used as an input for the transformer algorithm that computes the local attention for the whole sequence and the global attention for certain specific positions. After this step, the TE sequences that were not used for training are evaluated with the Transformer model and a TE class is predicted and compared with the labeled data. During the training a checkpoint is saved after each epoch, which allows the user to restore the training process from a previous step in case the training needs to be restarted or extended. In the final step, the trained model is validated on the test-set, which consists of unseen and unbiased comparable data.

For the classification of TEs, we use sliding windows to compute the k-mers of the input sequence and tokenize these data for use with the Transformer model. The model then outputs a prediction score, which is normalized with the Softmax function and gives the probability of the input sequence of belonging to any of the classes specified in the model.

### 2.3 Datasets

In the previous version of TEclass, the software permitted the classification of TEs into four categories, namely SINEs, LINEs, LTRs, and DNA transposons. To enhance the classification capabilities and allow a more in-depth categorization of TEs and by taking advantage of the larger datasets available today we built a classification model that assign TEs to sixteen different categories of TEs. These categories are based on the Wicker classification and include within the LTR order: Copia, Gypsy, Pao, and ERV; within the LINE order: L1/L2 and RTE; from the DNA subclass 1: Tc1/Mariner, hAT, Merlin, Transib, P, and Crypton; Helitrons from the DNA subclass 2 and SINEs (refer to Table 2).

**Table 2.**
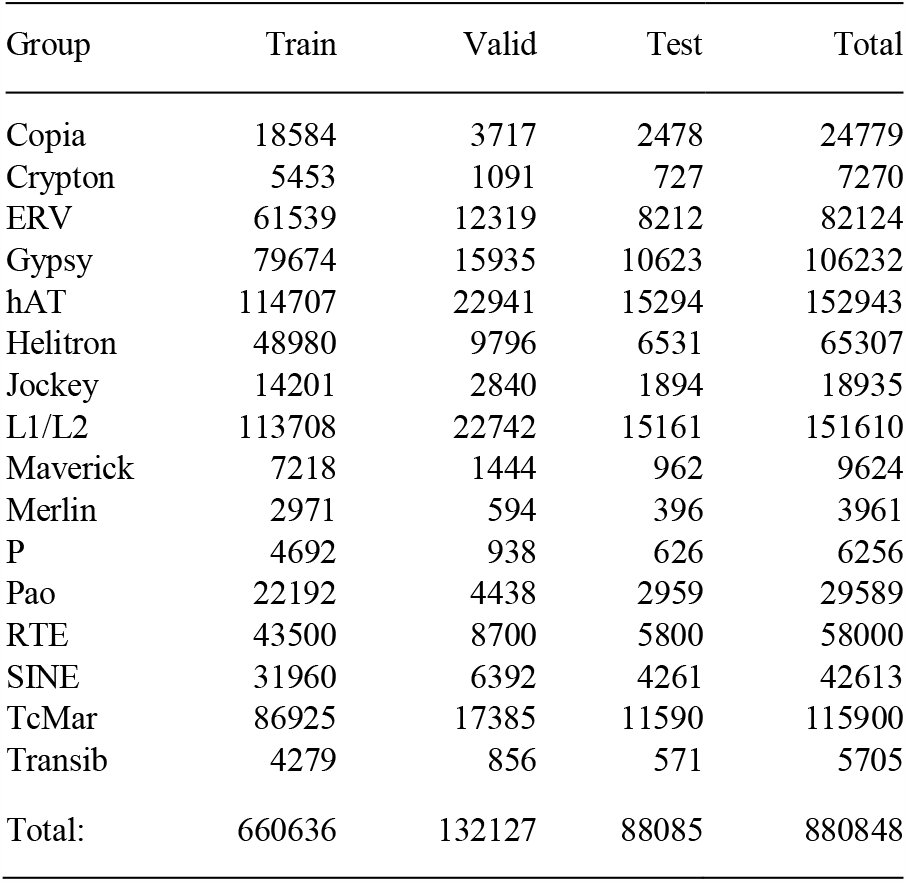
DFAM version 3.7 curated and non-curated data and Repbase version 18 used for training TEclass2, showing how the TEs from different categories were used for training, validation and testing.

| Group | Train | Valid | Test | Total |
| --- | --- | --- | --- | --- |
| Copia | 18584 | 3717 | 2478 | 24779 |
| Crypton | 5453 | 1091 | 727 | 7270 |
| ERV | 61539 | 12319 | 8212 | 82124 |
| Gypsy | 79674 | 15935 | 10623 | 106232 |
| hAT | 114707 | 22941 | 15294 | 152943 |
| Helitron | 48980 | 9796 | 6531 | 65307 |
| Jockey | 14201 | 2840 | 1894 | 18935 |
| L1/L2 | 113708 | 22742 | 15161 | 151610 |
| Maverick | 7218 | 1444 | 962 | 9624 |
| Merlin | 2971 | 594 | 396 | 3961 |
| P | 4692 | 938 | 626 | 6256 |
| Pao | 22192 | 4438 | 2959 | 29589 |
| RTE | 43500 | 8700 | 5800 | 58000 |
| SINE | 31960 | 6392 | 4261 | 42613 |
| TcMar | 86925 | 17385 | 11590 | 115900 |
| Transib | 4279 | 856 | 571 | 5705 |
| Total: | 660636 | 132127 | 88085 | 880848 |

For the creation of the curated dataset for training the models, we used TE sequences from Dfam, curated and non-curated version 3.7, and the curated Repbase version 18. In addition, the curated database was very limited and too small for models that are more complex. To allow more categories of TEs, we need a larger dataset with a larger variety of TE classes. The use of a non-curated database improves the number of data points, albeit with a slight decrease in the quality of the results.

From all the TE sequences in the dataset 75% are used for training the model, 15% are used for validation, and 10% for testing (Table 2).

### 2.4 Evaluation of TE models

In order to evaluate the performance of the trained models we calculated the precision, recall and F1 score of each one of the TE classes of a model to assess its reliability in the classification. Precision is a measurement of the exactness of the model and is the number of true positives divided by the sum of true positives and false positives. A high value of precision indicates that false positives are rare in the model. Recall is a measurement of the ability of the model to identify all the correct predictions and is defined as the proportion of true positives divided by the sum of true positives and false negatives. A high value of recall indicates a good level of completeness of the classification model, meaning that false negatives are not common. Although both measures are useful, they have their own limitations, as precision does not consider misclassifications meanwhile recall does not consider false positives. The F1 score combines both measures as it is the harmonic mean of precision and recall, where both contribute equally to the score, which has values between 0 and 1.

Since in each model we are using multiple classes with different number of TEs for comparing whole models between themselves, we calculated the F1 weighted average using the following formula:

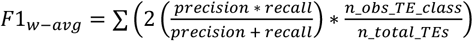

The weighted F1 score takes into consideration the class imbalance of the models and calculates the F1 score for each TE class, which is then multiplied by the proportion of TEs from that class in the total dataset. During the training of the models, these scores are calculated after each training cycle or epoch, as we used 10% from each TE class of the dataset for testing the performance of each model, as shown in the column “Test” from Table 2. In this way, during the training the trajectory of the changes in the performance can be checked in real-time. We also evaluated the performance of the software by classifying TEs models obtained from annotated genomes.

### 2.5 Software and hardware specifications

The hardware used for training were an Nvidia GTX 1660Ti 6 GB, RTX 3080 12 GB, A100 40 GB, and A100 80 GB. Palma II (https://confluence.uni-muenster.de/display/HPC), the HPC of the University of Münster, provided the latter two. These tests were conducted using float32 training, as mixed precision training is still marked as an experimental feature in the Trainer API. Gradient Accumulation was used for models based on non-curated data or very complex parameters, to overcome memory constraints. TEclass2 was developed using Python3 and requires NumPy (Harris *et al*., 2020), scikit-learn (Pedregosa *et al*., 2011), PyTorch (Paszke *et al*., 2019), Tensorboard (Abadi *et al*., 2016), Transformers, and Tokenizer packages (Wolf *et al*., 2020).

## 3 Results and Discussion

### 3.1 Parameters used for training TE models

During the training of the models, we evaluated different configurations, pre-processing steps, and parameters regarding the complexity of the models. We tested building models using 4-mer and 6-mer, but this resulted in a very similar performance to the default 5-mer models. As expected, allowing longer sequence inputs by increasing the embedding size provides an overall improvement of the results in particular for longer TEs. We noticed the importance of using dilated sliding windows to improve the results when the input contained long sequences. This resulted in an increased F1 score, which impacted the overall performance of the trained model.

Increasing the sparse attention did not improve the precision, indicating that local attention did not have an impact on classification beyond a specific minimal attention window and increasing it did not produce significant improvements. We also noticed that the complexity of each layer beyond a certain point did not increase the performance metrics, but what affected the most was the complexity of the model itself. In contrary to other methods of model complexity, the use of additional global attention slightly increased the overall scores.

To summarize, the complexity of the model and the available data points did not change the performance as much as expected. This introduced a grain fine-tuning with different parameters for TE-class2 (Table 3). The classifier approach using 5-mers with dilated sliding windows, using the maximum embedding size with 2048 tokens alongside a few additional global attention tokens achieved the highest scores.

**Table 3.** Optimal parameters used to train the Transformer model with both curated and non-curated dataset. These parameters may not be optimal for other datasets and their values depend on the hardware resources available.

| Parameters | Values | Description |
| --- | --- | --- |
| num_epochs | 100 | Times the model is trained on the same data. |
| max_embeddings | 2048 | Maximum length of the input sequence, but can be overcome with the Longformer model. |
| num_hidden_layers | 8 | Dimensions of the encoder hidden state for processing the input. |
| num_attention_heads | 8 | Number of heads (neurons) used to process input data and make predictions. |
| global_att_tokens | [0, 256, 512] | Positions of the input where all the sequence is considered |
| intermediate_size | 3078 | Dimensionality of the feed-forward layer |
| kmer_size | 5 | Length of the words for dividing the input sequence. |

#### 3.2 Training of TE models

We built a Transformer model that allows the classification of TEs in sixteen categories and the performance results with the training dataset can be observed from a confusion matrix (Figure 2) and a table of model testing statistics (Table 4),

**Table 4.** Evaluation parameters obtained from curated dataset using 16 TE classes.

| Group | Precision | Recall | F1-score | Support |
| --- | --- | --- | --- | --- |
| Copia | 0.69 | 0.60 | 0.64 | 3716 |
| Crypton | 0.62 | 0.69 | 0.65 | 1090 |
| ERV | 0.86 | 0.91 | 0.88 | 12318 |
| Gypsy | 0.81 | 0.71 | 0.76 | 15934 |
| hAT | 0.84 | 0.82 | 0.83 | 22941 |
| Helitron | 0.72 | 0.81 | 0.76 | 9796 |
| Jockey | 0.61 | 0.71 | 0.66 | 2840 |
| L1/L2 | 0.89 | 0.80 | 0.84 | 22741 |
| Maverick | 0.56 | 0.64 | 0.60 | 1443 |
| Merlin | 0.64 | 0.70 | 0.67 | 594 |
| P | 0.35 | 0.60 | 0.44 | 938 |
| Pao | 0.71 | 0.70 | 0.70 | 4438 |
| RTE | 0.76 | 0.78 | 0.77 | 8700 |
| SINE | 0.76 | 0.80 | 0.78 | 6391 |
| TcMar | 0.76 | 0.77 | 0.76 | 17385 |
| Transib | 0.67 | 0.75 | 0.71 | 855 |
| Accuracy: |  |  | 0.79 | 132120 |
| Macro avg: | 0.70 | 0.74 | 0.72 | 132120 |
| Weighted avg: | 0.79 | 0.79 | 0.79 | 132120 |

**Fig. 2.**
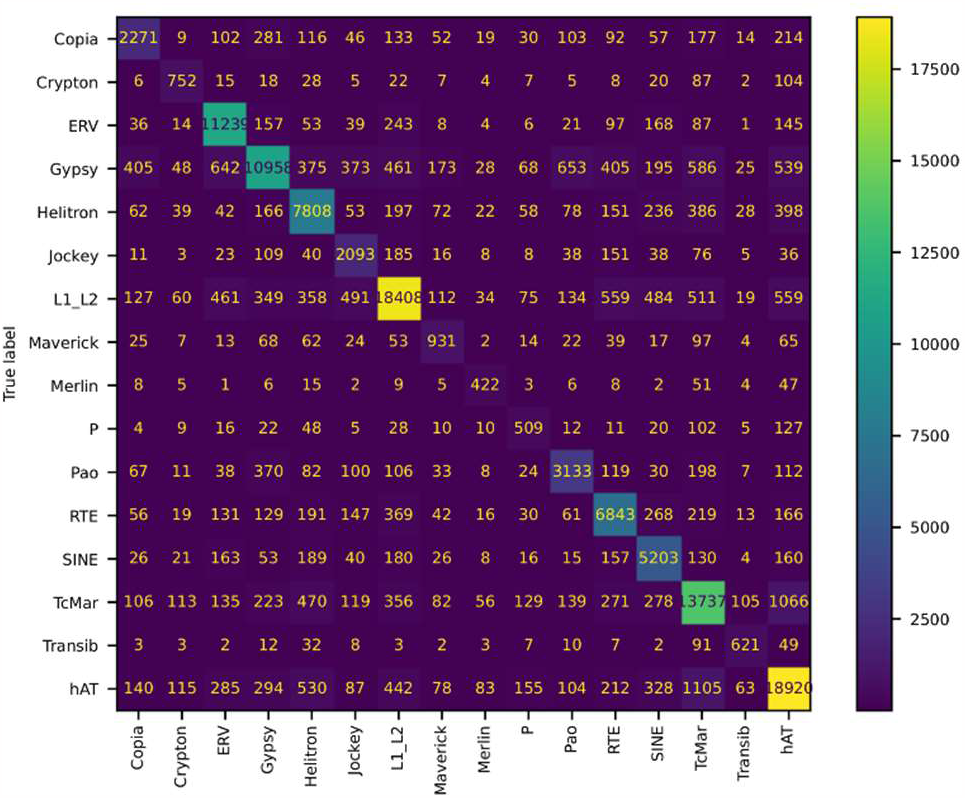
Confusion matrix for the training of non-curated data from Dfam version 3.7 and Repbase version 18.

For classifying TEs beyond just their main order, as the previous version of TEclass does, the data from curated databases was insufficient, so we relied on the Dfam version 3.7 non-curated dataset and Repbase version 18. We acknowledge the risks involved in using non-curated data, yet we expect that the majority of the sequences will be correctly classified, outweighing biases introduced by those erroneously labeled.

As expected, when more categories are added in order to train a model, the model increases in complexity, takes more time to train, and there is a greater chance of misidentification. Furthermore, some classes of TEs are similar to each other, making classification difficult. When we train a model using sixteen different classes of TEs, we observe good performance, considering the size of the model for the hAT, L1/L2, and ERV transposons, all obtaining F1-scores greater than 0.8. The performance of the model in classifying the TcMar, SINEs, RTE, Transib, Helitrons, and Gypsy transposons is also quite acceptable, with F1-scores for all of them greater than 0.7. The performance with some other transposons, like Copia, Crypton, Maverick, Merlin, and P, was suboptimal, with F1-scores below 0.7. In general, the weighted average of the model has an F1-score of 0.76, with a precision score of 0.77 and a recall of 0.76.

The misclassification that we observed can be attributed in part to the limitation of the Transformer model itself or to how the models were built, but can also be attributed to mislabeled data in the TEs dataset or to the lack of enough representative sequences for training the model in a given class of TEs.

### 3.3 Comparison with other machine learning tools

To assess the classification performance of the TE model trained with TEclass2, we employed TE models built using RepeatModeler2 (Flynn *et al*., 2019) version 2.0.3 with default parameters and the LTR-detector module on the fruit fly and zebrafish genomes, both obtained from the UCSC Genome Browser database.

RepeatModeler already provides TE model classifications through its RepeatClassifier module. To further curate these results, we ran nhmmscan from the HMMER package version 3.3.2 (hmmer.org), with the parameters “--noali -E 1e-10” to align these TE models with the curated Dfam database version 18. Sequences that exhibited poor alignment with TEs or had hits to TEs from different orders were discarded, as this often indicates chimeric models, a common problem in most TE model building software (Rodriguez and Makałowski, 2022).

We ran the TE models obtained from the rice and fruit fly genome with TEclass2 and compared them to the classification assigned by RepeatModeler. We also ran these TE models using the default settings of two other machine learning software, DeepTE and TERL.

Making a direct comparison of TE classification tools can be difficult, as some of them specialize in solving specific problems, such as differentiating TEs within the same class or classifying into a different number of families. For the 75 TE models built from the rice genome, most of them are Gypsy and Copia, but are also some Helitrons and LINEs. TEclass2 using a minimum probability threshold of 0.7 was able to classify 79% of the TEs models and from those 82% were identified correctly at superfamily level and for 87% the order was also predicted accurately.

For the fruit-fly genome, 667 TE models were built, with a diverse number of TE models from Gypsy, Jockey, Bel-Pao, R1, Tc1-Mariner, Copia, P, Helitron, Transib, and hAT. From this dataset, 393 were classified with a probability of at least 0.7, that is 59% of the total, and from those 285 were correctly classified, meaning a 73% accuracy (see Supplementary Material).

Adopting a stricter probability threshold, such as 0.9, improved the TE prediction accuracy to 83%, but reduced the number of TEs that are classified, to 281 out of 667. During these analysis there were also a small group of TEs that couldn’t be classified because they don’t belong to any of the families included in the trained model.

For testing other machine learning tools, we ran DeepTE with the parameters “-m M -sp M” for the fruit fly TEs models in order to specify that TEs are from a Metazoa genome and “-m P -sp P” for rice TEs models to specify that the TEs are coming from a plant genome. DeepTE sets by default a limit of a probability of 0.6 to give a result, if the value is lower, then the TE is not classified.

For the TEs models built from the rice genome, the accuracy for predicting the TE order was of 53%, and for the superfamily of 45%. From all the TEs models built, only 79% had their order predicted and 29% their superfamily. For the TE models built from the fruit fly genome, DeepTE was able to predict the order accurately for 35% and the superfamily for 32%. The proportion of TEs with the order predicted was 88% and for the superfamily 58% (See supplementary material).

We also ran TERL for both genomes, using the model DS3, which is a model that can classify TEs into twelve superfamilies. In the case of TE models constructed from the rice genome, the accuracy of predicting the TE order was 68%, and 50% for the superfamily. Among all of the TE models constructed, a classification rate of 91% was achieved as 9% were classified as non-TEs.

For TE models from the fruit fly genome, 42% were accurately predicted and 21% for the superfamily. Approximately 57% of the TEs had their order predicted (See supplementary material).

### 3.4 TEclass2 web interface

The models built with curated and non-curated data can be used to classify TEs at:

https://bioinformatics.uni-muenster.de/tools/teclass2/index.pl

The web interface provides a simple interface that allows the classification of TEs models DNA sequences into sixteen superfamilies and the possibility to filter the results by a probability value. The TE sequences used as input can be uploaded as a FASTA file or inputted in the text box.

The output produced by the classification module provides an accessible and compact format as a table or as a downloadable TSV file useful for further analysis. This table includes the classification predicted, with the order and superfamily that corresponds to the class with the highest probability, and the Softmax confidence scores for each class in the model (Figure 3). For training models it is necessary to download the source code of the program from https://github.com/IOB-Muenster/TEclass2 and follow the configuration instructions in order to run it.

**Fig. 3.**
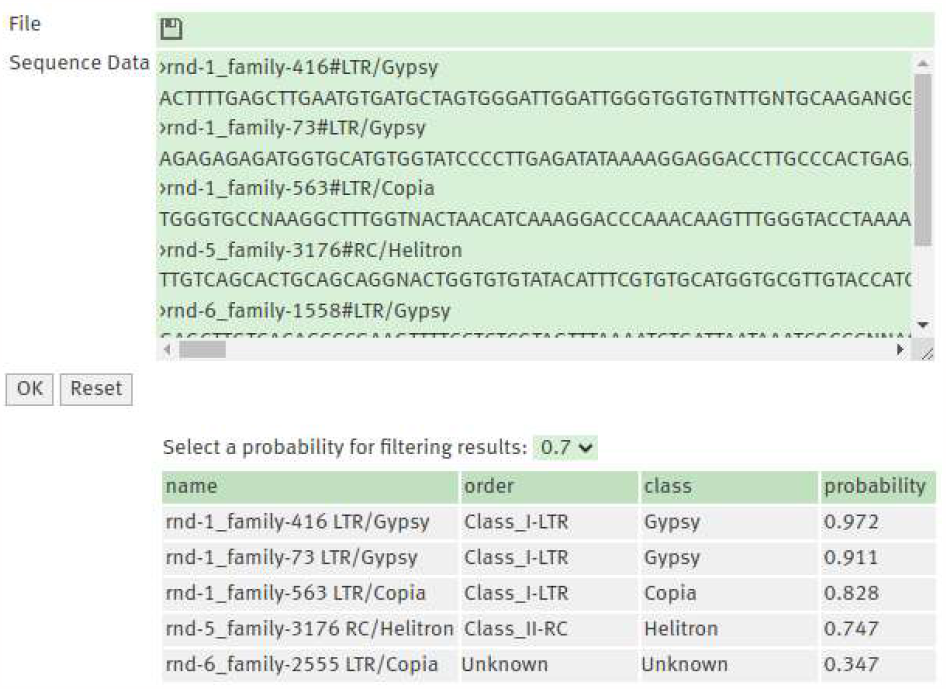
The web interface for TEclass2 allows inputting the TE sequence to be classified in a text box or as a FASTA file. The output shows the classification and the Softmax values the model scored for each class of TE.

### 3.5 Known limitations

Some of the limitations of TEclass2 are the hardware requirements for training new models, as it is necessary to have a GPU to build new models. Other weaknesses come directly from the Transformers architecture (Dong *et al*., 2021), as the training of models require a fine-tuning of multiple parameters, which may lead to several trial and error cycles. Inherent to the architecture comes the difficulty to control the attention mechanism, which can put attention in parts of the sequences that are biologically not relevant or show an undesirable attention-bias. The algorithm also needs to fragment the sequences into k-mers meaning that the sequence is not analyzed as a whole but divided into words similarly to NLP inputs.

Another issue may come from the input data itself, as machinelearning strategies require a large number of TE sequences that are well curated. This may be impossible to attain for some classes of TEs where only a few copies are present in curated databases.

## 4 Conclusions

TEclass2 is a comprehensive classifier that applies a state-of-theart Transformers architecture and yields promising results. When we trained many TE models, we computed a weighted overall accuracy of 79% for properly classified sequences when dividing TEs into sixteen families. TEclass2 is readily accessible for downloading or online use, providing researchers with a simple and accessible software for classifying unknown TE models.

## Acknowledgements

This work utilized the computational resources of the PALMA-II HPC cluster of the University of Münster (https://confluence.unimuenster.de/display/HPC)

